# A Smart Bait Dispenser with Real-time Monitoring for Species-Specific Bait Delivery

**DOI:** 10.64898/2026.10.25.761189

**Authors:** Prithul Chaturvedi, William M. Connelly, Tim Lennard, Chrissie Ee Been Ong, Timothy J. Smyser, Andrew S. Flies

**Author notes:** Corresponding author: Andrew S. Flies.

## Abstract

1. Remote image capture using trail cameras has become an essential part of many wildlife monitoring and management programs. Oral bait vaccination has been a valuable tool in wildlife disease management, but baits are generally dropped on the ground or delivered by devices that are typically developed to utilise physical and behavioural characteristics of the target species. Both approaches can result in a high proportion of baits being consumed by non-target species. Merging years of trail camera research knowledge with new technologies allows for development of more specific bait delivery options.
2. We developed an open-source smart bait dispenser that uses field-based image classification to selectively deliver baits to the target species. The system’s long-range wide area network-based remote telemetry provides updates to an online dashboard in near real time to show camera triggers, target species detections, and baits dispensed. We trialled four dispensers with placebo baits for 62 nights across three field sites in Tasmania, Australia to deliver baits to Tasmanian devil (*Sarcophilus harrisii*).
3. The dispensers recorded 2106 motion triggers, of which 876 were triggered by animals. The onboard devil classification model achieved 98.6% accuracy, 75.0% precision, 77.4% recall, and a 76.2% F1-score. The dispenser released 41 placebo baits in total, of which 31 were target-specific distributions in response to presence of Tasmanian devils at the dispenser, hence achieving an overall bait delivery success rate of 75.6%. Importantly, no animal was able to forcibly retrieve baits from the dispenser. Observed average battery life across trials was 19.2 nights.
4. These results clearly demonstrate the effectiveness of our innovative bait dispenser for targeted bait delivery, offering a versatile ecological tool that can be tailored for various species by simply updating the image classification model. This adaptable system can support critical initiatives like wildlife oral bait vaccination and supplementary feeding after wildlife translocations.

## Introduction

Oral bait vaccines (OBVs) have been an effective wildlife disease management strategy for five decades across five continents (Rupprecht et al., 2024; Steck et al., 1982). For a multi-species pathogens like rabies, vaccine-laden baits can be distributed either by hand in urban settings or dropped from aircrafts for broad-scale distribution. With the success of the rabies control, OBVs have been suggested or trialled for other species and diseases, such as the devil facial tumour disease (DFTD) in Tasmanian devils (*Sarcophilus harrisii*) (Flies et al., 2020), chronic wasting disease and bovine tuberculosis in cervids (Zimmerling and Napper (2026), reviewed in VerCauteren and Lavelle (2026)), and Sylvatic plague in prairie dogs (*Cynomys spp.)* (Rocke et al., 2017).

The effectiveness of OBV programs for wildlife hinges on vaccine baits reaching the target species. Consumption of baits by non-target species reduces vaccine availability for the target species and wastes limited funds by vaccinating the wrong species. Mechanised bait dispensers have been shown to improve target-specific bait uptake (Bastille-Rousseau et al., 2024; Bjorklund et al., 2017; Boulanger et al., 2006; Dempsey et al., 2022; Smyser et al., 2015b). However, dispensers are often designed to exploit the physical or foraging behaviour of the target species to maximise target-species specificity. Such a species-specific approach is preferred in poison-based pest management, where exclusion of non-target species is crucial, for example with invasive mustelids (*Mustela spp.*) (King et al., 2007) and for feral pigs (*Sus scrofa*) (Lavelle et al., 2018).

Bait dispensers can be designed out of common materials available at local hardware stores. A polyvinyl chloride (PVC) pipe-based bait station has been trialled in rabies vaccination of raccoons (*Procyon lotor*) in suburban United States (Boulanger et al., 2006). This dispenser was also modified to treat raccoons for roundworm (*Baylisascaris procyonis*) (Smyser et al., 2015a; Smyser et al., 2013). During trials of bait consumption by raccoons, baits distributed freely into the environment (i.e., ground baiting) were rapidly consumed (90% consumed within the first three days), thus limiting the temporal window for vaccination. However, the use of a bait dispenser with an automated timer for bait delivery successfully prolonged bait availability (Smyser et al., 2015a). When this dispenser was adapted for Tasmanian devils, it improved target bait uptake by 10-fold compared to ground baiting (Dempsey et al., 2022). However, the dispenser did not prevent eastern quolls (*Dasyurus viverrinus*) and common brushtail possums (*Trichosurus vulpecula*) from removing baits. Additionally, due to their smaller home range (Andersen et al., 2020; Jones & Rose, 2001), these animals frequently visited the dispenser and reduced bait availability for the target species. DFTD has led to low devil population density across the island state of Tasmania, reducing the probability that devils will find baits before competitors. An improved dispenser to target endangered species like the Tasmanian devil is thus required (Dempsey et al., 2022).

Recent developments in wildlife management have leveraged technology to identify the target species. The Felixer Grooming “Trap” uses infrared sensors to identify and control invasive red foxes (*Vulpes vulpes*) and feral cats (*Felis catus*) by estimating the animal’s dimensions and spraying a dose of poison onto the fur of the identified target species (Moseby et al., 2020). A new version of the device uses a camera and an artificial intelligence model in species identification (Thylation, 2026). However, the bulky structure and heavy weight of the Felixer require the system to be airlifted to remote locations (Department of Lands Planning and Environment, 2025). Additionally, this approach is not well-suited to vaccine delivery.

A five-bait smart dispenser was successfully trialled in Australia for wild dogs (*Canis familiaris*) and foxes (Charlton et al., 2023) using Raspberry Pi 3 Model B+ to run a deep learning model for species identification and targeted bait delivery. However, due to high power consumption of the chosen single board computer, this device is unsuitable for multi-week operation on a single charged battery. The authors also called for design improvements to accommodate more baits and reduce mechanical noise. A study on edge devices determined the ArduinoPro Portenta H7 Vision Shield (LoRa) (hereby Portenta) and ATtiny85 microcontrollers were better suited for an artificial intelligence camera-based bait dispenser because of its power efficiency and computational capabilities (Chaturvedi et al., 2026).

With the goal of designing a generalisable, scalable, cost-effective, and open-source ecological tool, we developed a camera-based smart bait dispenser that uses image classification to identify the target species and deliver baits *in situ*. We implemented the dispenser with placebo baits for Tasmanian devils over 62 nights at three sites in Tasmania. We used a simple convolutional neural network (CNN) model to classify devils from non-target species. We assessed the model’s performance using accuracy, precision, recall, and F1-score. We also used the Portenta’s long range wide area network (LoRaWAN) connectivity to receive activity updates on an online dashboard in near real-time. Paired trail cameras deployed with dispensers were used to evaluate animals’ interactions with the dispenser and provide an independent platform to quantify species visitation rates. We determined the dispenser’s overall success by its ability to accurately dispense bait for the target species and its feasibility for multi-day field deployment.

## Methods

### 2.1 Design

#### 2.1.1 Dispenser shell and carousel design

**Fig. 1** illustrates the design of the bait dispenser. The outer shell was constructed from a Ø250 × 80 millimetre (mm) polyvinyl chloride pressure pipe. A 50 × 50 mm square-shaped opening was cut on the underside of the shell to allow gravity-based bait drops. The front and rear ends of the shell were enclosed with laser cut transparent acrylic discs that were 6 mm thick. The rear acrylic sheet was permanently glued to the shell using cyanoacrylate adhesive (super glue).

The dispenser comprises four main internal components: (i) a central hub fixed to the rear acrylic disc; (ii) a carousel, capable of holding up to 20 baits, that rotates around the hub driven by a 5-volt geared direct current motor and uses angled serrations to determine the stopping position based on a roller limit switch; (iii) a sleeve that houses the battery and supports the system’s printed circuit board; and (iv) a mount for the infrared (IR) lights. The IR lights were positioned above the motor. The printed circuit board was designed using Fritzing (Knörig et al., 2009) and ordered from PCBWay (China).

#### 2.1.2 Electronic components

The bait dispenser was powered by a generic 14.4 V 4 series 1 parallel (4S1P) 18650 battery pack. The batteries were charged prior to use using IMAX B6 LiPro Balance Charger (Seeed Studio, 2024). Two SY8205-based buck converters supplied regulated 5 V or 12 V power to the dispenser’s electronic modules (DFRobot). An ArduinoPro Portenta H7 Vision Shield (LoRa) (Arduino, 2020a, 2020b) equipped with a Himax HM0360 NoIR camera (Himax, 2020) was used as the primary image capture, computational, and communications device. Note that commercially available HM0360 cameras are equipped with an infrared-cut filter that must be manually removed to enable nighttime imaging under infrared illumination. However, this procedure may damage the lens or image sensor. HM0360 cameras without an infrared-cut filter (NoIR versions) can be obtained by contacting Arducam. An ATtiny85 (Atmel, 2013) was used to manage deep sleep and reduce power consumption, as Portenta’s native sleep modes do not adequately reduce energy consumption for long-term deployment (Chaturvedi et al., 2026). Motion detection was achieved using an HC-SR501 passive infrared (PIR) sensor, connected to both the Portenta and ATtiny85. A DS1307-based real time clock module (Analog Devices, 2015) was used to maintain accurate timestamps on dispenser activities. A DRV8871 motor driver module (Texas Instruments, 2016) controlled the carousel motor.

**Figure 1.**
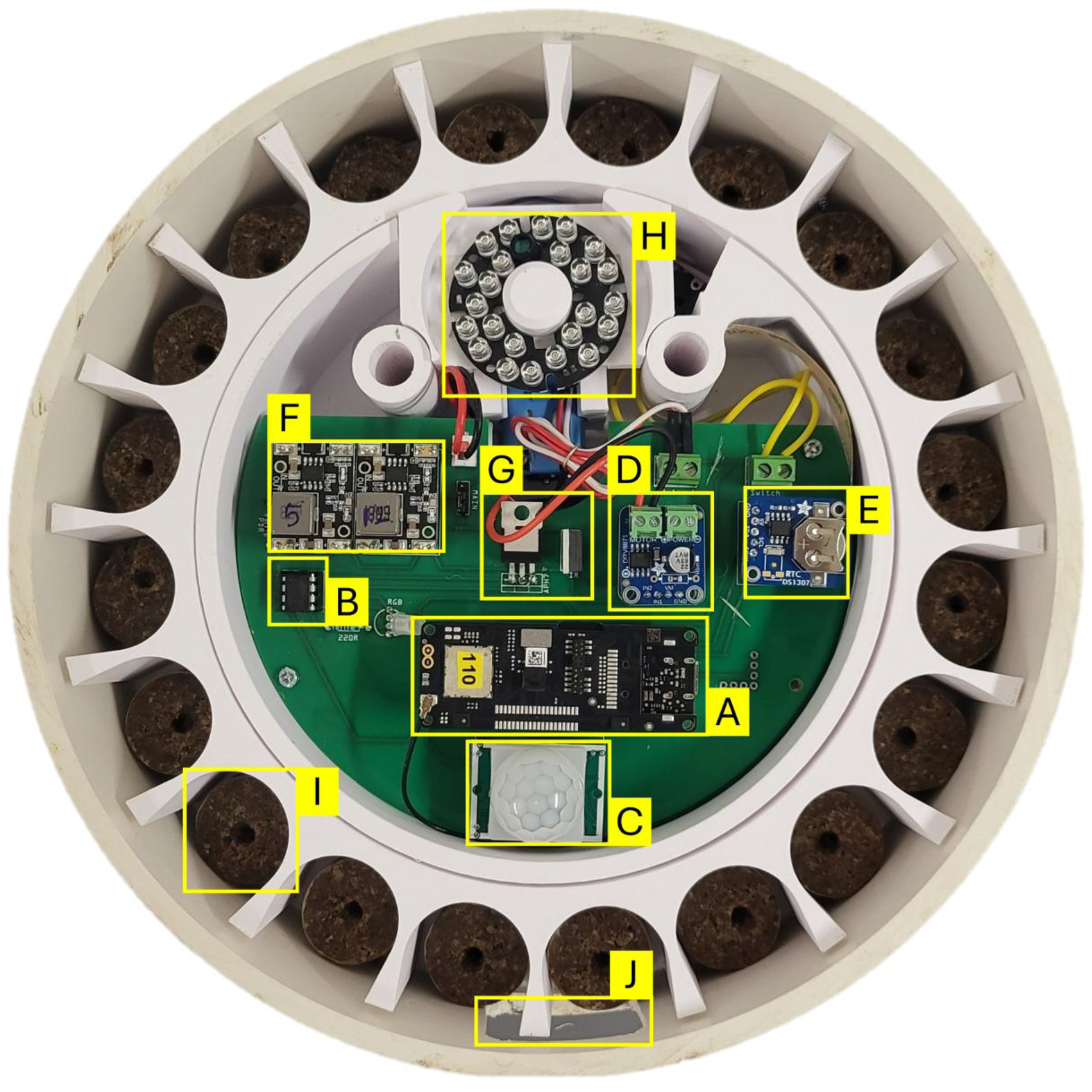
Anatomy of the smart bait dispenser. (A) Arduino Portenta H7 Vision Shield (LoRa) with device identity label “110”; (B) ATtiny85; (C) HC-SR501 passive infrared sensor; (D) DRV8871 motor driver breakout board; (E) DS1307 real time clock breakout board; (F) SY8205 buck converter breakout boards; (G) IRF4905 and IRLB8721 MOSFETs; (H) Infrared illuminator; (I) FOXOFF® placebo bait; (J) A square window to release baits for the target species. The 5V DC motor is placed behind the infrared illuminator. The printed circuit board is placed on top of a 3D printed sleeve that holds the 14.4V 4S1P 18650 battery pack.

### 2.2 Software

Custom firmware based on OpenMV and Ardunio frameworks were developed for the Portenta and the ATtiny85, respectively. Locally on the Portenta, we ran CNN1, an 8-bit integer-quantised convolutional neural network (Chaturvedi et al., 2026), to distinguish Tasmanian devils from non-target species. This binary classification model was trained on high-definition trail camera images of Tasmanian wildlife. CNN1 was selected owing to its high devil classification metrics (approximately 96%) and rapid inferential speed on Portenta (0.149 seconds). A classification threshold probability of 0.5 was applied for field studies, and a bait was released only when at least five of ten captured images were classified as devil. The threshold and number of image matches can be easily modified based on performance.

We considered four metrics to evaluate the model’s performance in our bait dispenser: accuracy, precision, recall, and F_1_-score. If F_P_, F_N_, T_P_, and T_N_ denote false positive, false negative, true positive, and true negative, respectively, then:

a. Confusion matrix:

|  |  | Predicted class |  |
| --- | --- | --- | --- |
|  |  | Target species | Non-target species |
| Actual class | Target species | T<sub>P</sub> | F<sub>N</sub> |
|  | Non-target species | F<sub>P</sub> | T<sub>N</sub> |

b. Accuracy = 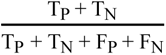
c. Precision = 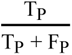
d. Recall = 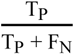
e. F_1_-score = 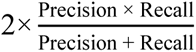

### 2.3 Operational workflow

The operational workflow of the smart bait dispenser is shown in **Figs. 2** (simplified flowchart) **and S1** (detailed flowchart). When motion is detected while the system is in deep sleep, the ATtiny85 powers the Portenta by high-side switching through an IRF4905 metal oxide semiconductor field effect transistor (MOSFET). The Portenta then executes all the functions of the bait dispenser. Each motion triggers the IR lights via low side switching through a IRLB8721 MOSFET and prompts the capture of 10 images with a 500 millisecond pause between images. These images are fed into the CNN1 model to classify the detected species and are simultaneously stored on the Portenta’s memory card. If CNN1 predicts the presence of the target species at the given threshold and number of matched images, the DC motor is gradually powered using the pulse-width modulation-based motor driver to rotate the carousel. Carousel positioning is determined by a roller limit switch fitted on the hub. Upon engagement of the switch, the motor stops, aligning the next bait with the release opening, allowing gravity to pull the bait out of the dispenser. Following bait release, the timestamp of the event is saved in a log file.

To reduce the likelihood of dispensing multiple baits to the same individual, the dispenser was set to enforce a configurable 40-minute lockout period during which no additional baits would be released. The dispenser will continue to collect images upon motion triggers during the lockout period. After two minutes (modifiable based on requirement) of inactivity, i.e., no new motion trigger, the Portenta prompts the ATtiny85 to switch off the IRF4905 MOSFET, hence returning the system to deep sleep mode. Before this shutdown, a message on the number of motion triggers, target species detections, and baits dispensed is saved to a log file. This information is also sent to our online dashboard made with Datacake (datacake.co/) via long-range wide area network (LoRaWAN). This process involves transmission of the message packet from the dispenser to the nearest LoRaWAN gateway, which is relayed to The Things Network (thethingsnetwork.org/) server via the internet (**Fig. 4 (d)**). Gateways in remote field locations need to be placed at high elevations so that its mobile data SIM card can connect to mobile network services.

**Figure 2.**
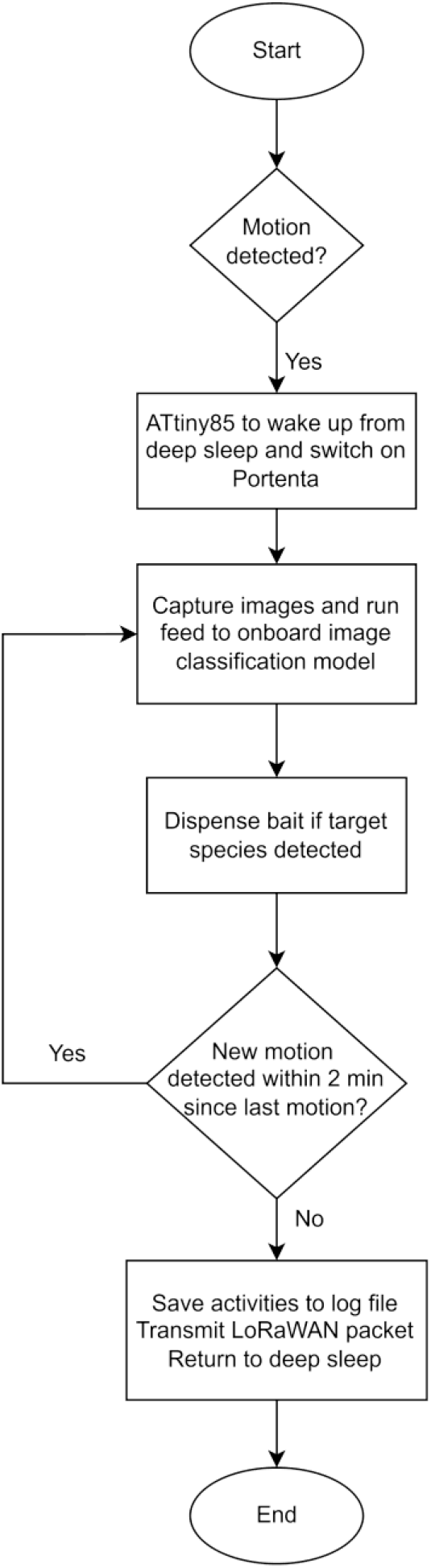
Flowchart of the bait dispenser’s operational workflow process. Made with draw.io (Jgraph, 2021).

### 2.4 Field trials

Trials were conducted at private properties in Margate, Crabtree, and Dunalley, Tasmania (**Fig. (a)**) where a diversity of species including devils, quolls, possums, and macropods (Bennet’s wallaby (*Notamacropus rufogriseus*) and Tasmanian pademelon (*Thylogale billardierii*)) had previously been observed. The Dunalley site, owned by Bangor Vineyard Shed, is located within a disease-free devil population conservation reserve and likely has the highest devil density in Tasmania. Hence, trialling here was strategic in achieving sufficient devil visitations. We deployed one dispenser each at the Margate and Crabtree sites and two at the Dunalley site. Collected data was divided into nightly periods, with each night defined as the 24-hour period from 12 PM to 12 PM the following day.

Each dispenser was loaded with 19 FOXOFF® placebo baits (Ø25 mm × 20 mm) (Animal Control Technologies Australia). A Reconyx HyperFire 2 trail camera and an 8-channel RAK7289 outdoor LoRaWAN gateway were also setup at each dispenser site (**Fig. 3 (b) and**). We used BotRf, a free Telegram Bot, to determine suitable locations to setup LoRaWAN gateways based on connection feasibility prior to field trials (Zennaro et al., 2017). The dispensers were mounted approximately 500 mm above ground using steel clamps attached to garden stakes. Fish oil was applied two-metres in front of the dispenser to attract animals to the optimal spot for image capture and classification. Using fish oil as a scent lure was used to replicate the trail camera setup from which the training data for the devil classifier was sourced.

As this study was conducted in the southern hemisphere, all the dispensers were oriented south to minimise false triggers caused due to sunshine and shadows movements. At the Margate and Crabtree sites, dispensers were inspected daily during the first four days of the trial to retrieve data from the dispensers and trail cameras, and perform maintenance as required. Dispensers at the Dunalley site were only remotely monitored and attended to when the dashboard indicated inactivity for >24 hours. Use of our bait dispenser with placebo baits in was approved by the University of Tasmania Animal Ethics Committee under #29566. No permit was required to access the trial sites.

**Figure 3.**
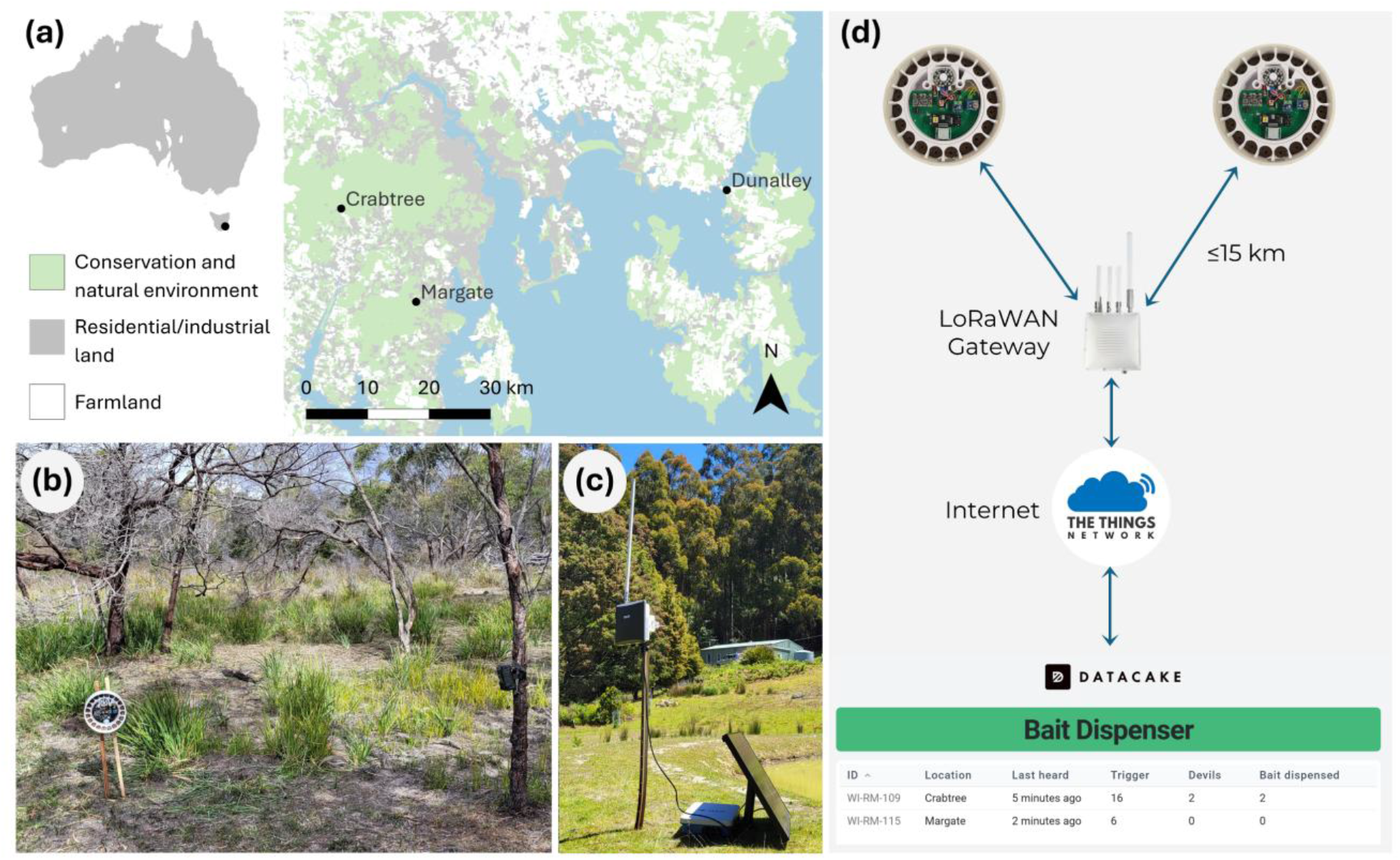
Field trial location and setup. (a) Dispenser trial sites in Tasmania, Australia. “NCH Land Use 2021” from theLIST ©StateofTasmania. Made with QGIS (QGIS Development Team, 2026). (b) A dispenser attached to garden stakes using steel clamps and a trail camera mounted on a tree; (c) RAK7289 LoRaWAN gateway setup with associated battery and solar panel was setup within 1 kilometre of the dispenser. (d) LoRaWAN network diagram. The bait dispensers send activity updates as message packets to the nearest gateway placed in an internet-rich location. The gateway uploads the packet to TheThingsNetwork server, which is then displayed on the online dashboard

## Results

### 3.1 Images captured on dispenser

**Table 1** provides a summary of all the motion triggers observed across the three sites. Triggers caused by animal visitation were confirmed using pictures captured on the dispenser. When the animal was not in the dispenser’s field of view (i.e., not captured on the dispenser’s camera), we used videos from the paired trail camera to identify the visitor by matching the activity date and time from devices. Validation with an independent device was crucial for assessing the dispenser’s motion-detection performance, since minimising false positives are important for optimal battery life and bait distribution. Cases in which neither the integrated bait dispenser camera not the independent trail camera recorded an animal were considered as false triggers.

**Figs. 4 and 5** show images of visiting animals captured by the dispenser’s Portenta camera. During the 62 trial nights, the dispenser image capture was triggered 2106 times, 876 (42.0%) of which were due to the presence of an animal. The remaining 1230 (58.4%) triggers were false positives, caused by environmental factors such as sunlight, shadows, and moving branches and grass that set off the motion detector.

Of the 876 motion triggers caused by animals, the dispenser captured an animal on 418 (47.7%) of those for the onboard model to classify the species. Across all sites, quolls were responsible for the greatest number of triggers (254, 29.0%), followed by macropods (241, 27.5%) and devils (198, 22.6%). Within sites, Crabtree had the highest motion triggers (368, 42.0%), followed by Dunalley (280, 32.0%) and Margate (228, 26.0%). The most observed species at each site were quoll at Margate (123, 53.9%), macropods at Crabtree (150, 40.8%), and devil at Dunalley (169, 60.4%). We observed 62 (14.8%) devils on the dispenser cameras. Out of these, only 4 (6.5%) were detected from Margate, 18 (29.0%) from Crabtree, and 40 (64.5%) from Dunalley.

### 3.2 Identifying the target species

The confusion matrix presenting the devil classification model’s predictions against actual

visitations is shown in **Table 2**. Corresponding performance metrics are as follows:

a. Accuracy: 98.6%
b. Precision: 75.0%
c. Recall: 77.4%
d. F1-score: 76.2%

The model’s strong ability to identify true negatives drives the high accuracy achieved. The model’s capability of recognising the target species is best explained by recall (ratio of correctly predicted devils out of all the truly observed devils).

### 3.3 Dispensing baits to the target species

**Table 3** presents data on all the baits dispensed across the three study sites. Including the 40-minute lockout period after dispensing a bait, the devil classification model initiated bait distribution 64 times (also indicated by the prediction class in **Table 2**). The four dispensers released 41 baits in total: 31 (75.6%) for devils and 10 (24.4%) for non-target species. At the Margate site, of the 2 baits released, both were triggered by the presence of devils (100% success). At Crabtree, 11 baits were released correctly for devils, and 8 were incorrectly released for non-target species (57.9% success). At Dunalley, 20 baits were released from the two dispensers, of which 18 were correctly dispensed to devils (90% success). Based on the log files, we conclude that the dispensers failed to release baits 8 times due to potential mechanical issues or insufficient power to the motor due to low battery power. The dispensers withheld baits 15 times during the lockout period. Importantly, the trail cameras recorded videos of the dispenser functioning (https://youtu.be/hayhBW1BI3k) and devils consuming baits (https://youtu.be/3etGtvsIoOc). Note that in the video of the devil consuming the bait, we merge three trail camera videos into a single video to show a more complete bait consumption session. The trail cameras also caught several quolls, possums, and devils trying to forcibly retrieve baits by biting the dispenser and inserting their snouts and forelimbs into the bait-release opening to turn the carousel (**Fig. 6 (c) to (f)**). It was observed that no animal could succeed in stealing baits from the dispenser unless the dispenser initiated a bait release.

### 3.4 Battery life

A primary concern before field trials was battery life. The bait dispenser at the Margate site was active for 12 nights, after which the battery pack was replaced. After this, the dispenser lasted 13 nights in the second run. The dispenser at the Crabtree site was active for 7 nights. The two dispensers at the Dunalley site were active for 31 and 33 nights. We observed numerous false-positive motion triggers at the Margate and Crabtree sites, resulting in higher computational activity and, consequently, shorter battery life than at the dispensers the Dunalley site.

**Figure 4.**
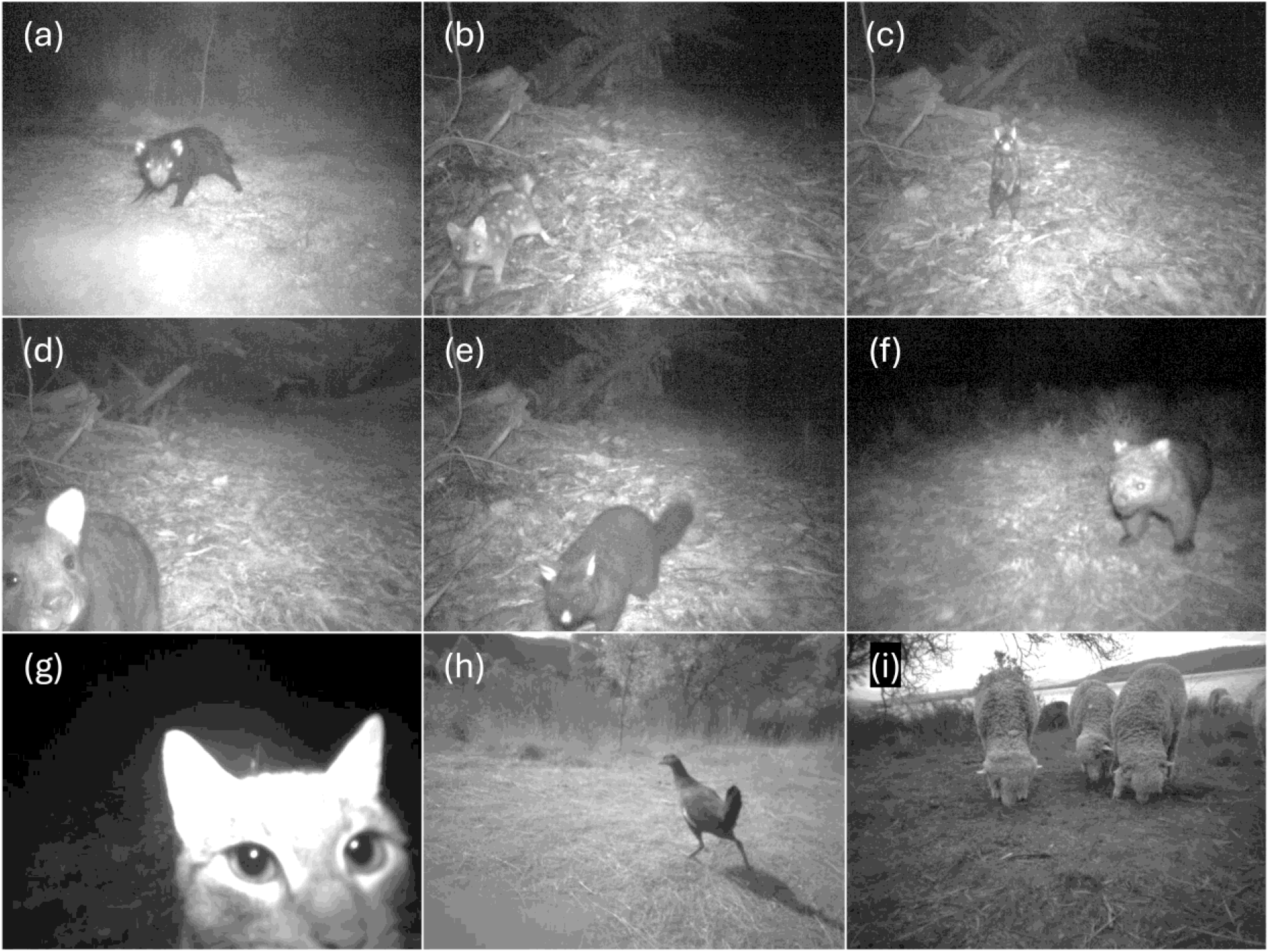
Images captured on the dispenser’s Himax HM0360 NoIR camera. (a) Tasmanian devil; (b) tan morph Eastern quoll; (c) black morph Eastern quoll, (d) Bennett’s wallaby; (e) common brushtail possum; (f) common wombat (*Vombatus ursinus*); (g) cat (*Felis catus*); (h) Tasmanian native hen (*Tribonyx mortierii*); (i) sheep (*Ovis aries*).

**Figure 5.**
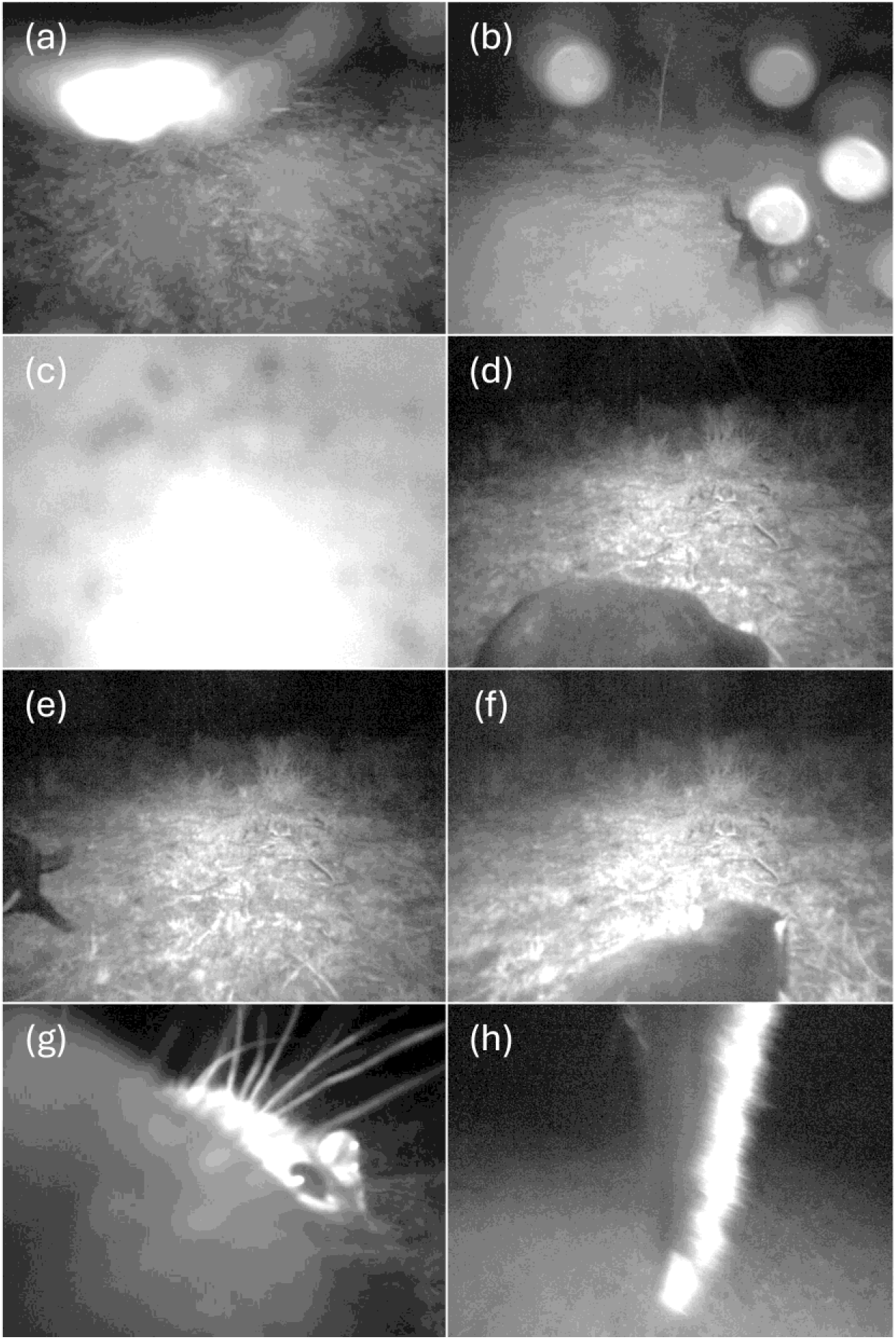
Images captured on the dispenser where animals are not in the field of view. (a) dispenser falsely triggered during rain; (b) image of a devil distorted by a raindrop in front of the camera lens; (c) image clicked when frost was set on the front acrylic disc; (d) a devil too close to the dispenser with no identifiable feature visible for a successful image classification. (e) a devil moved out of frame; (f) backside of a devil which can be mistaken for a possum or a wombat; (g) a devil very close to the dispenser; (h) tail of a quoll that climbed on top of the dispenser.

**Figure 6.**
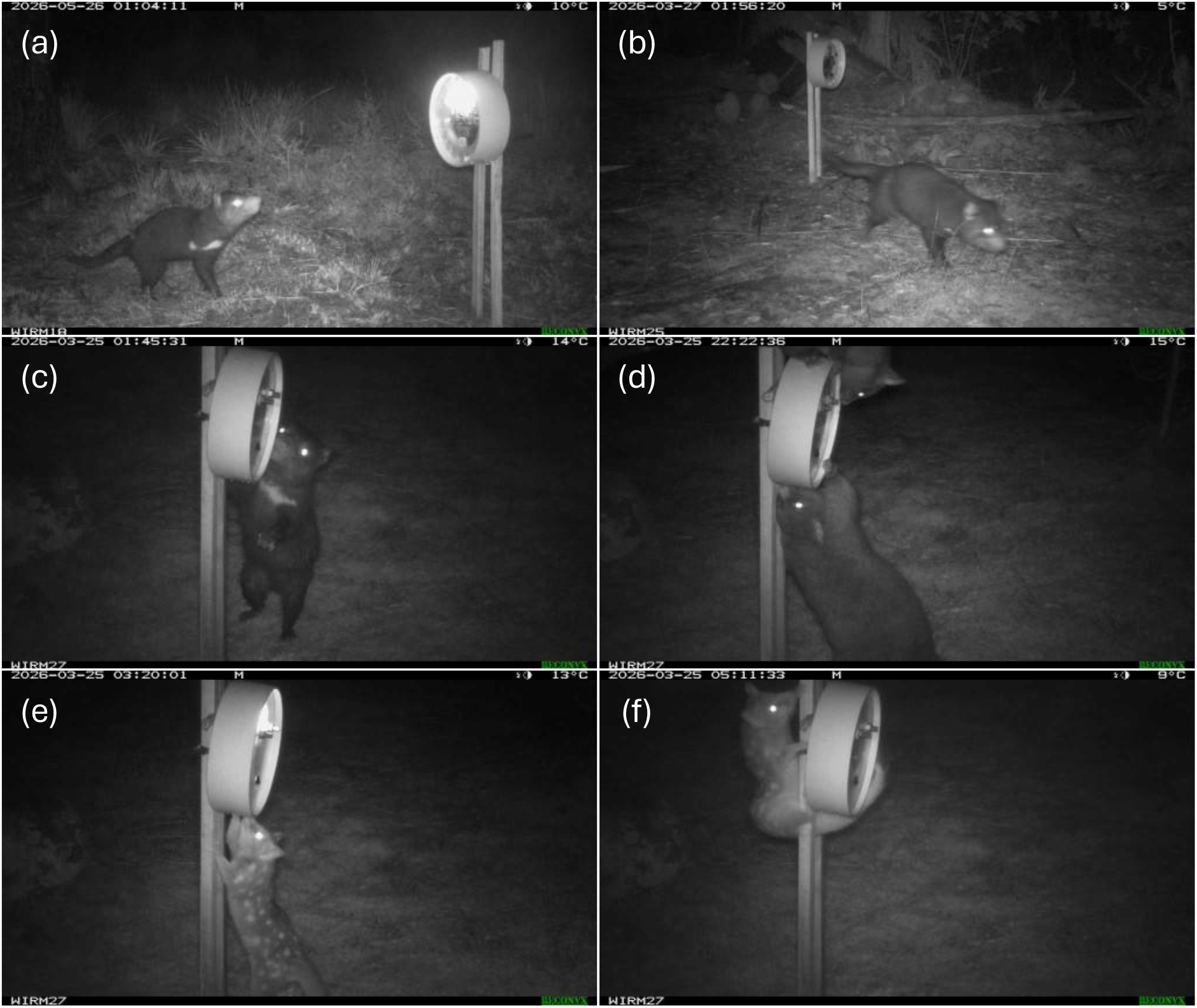
Animal interactions with the bait dispenser. (a) a Tasmanian devil in the ideal field of view of the dispenser; (b) a devil quickly moving past the dispenser without triggering it; (c) a devil lured by the scent of baits in the dispenser, trying to turn the carousel with its right forelimb; (d) two possums trying to remove baits by force; (e) a quoll trying to turn the carousel of a frosted dispensed to drop the bait; (e) the same quoll trying another approach to remove baits.

**Table 1.**
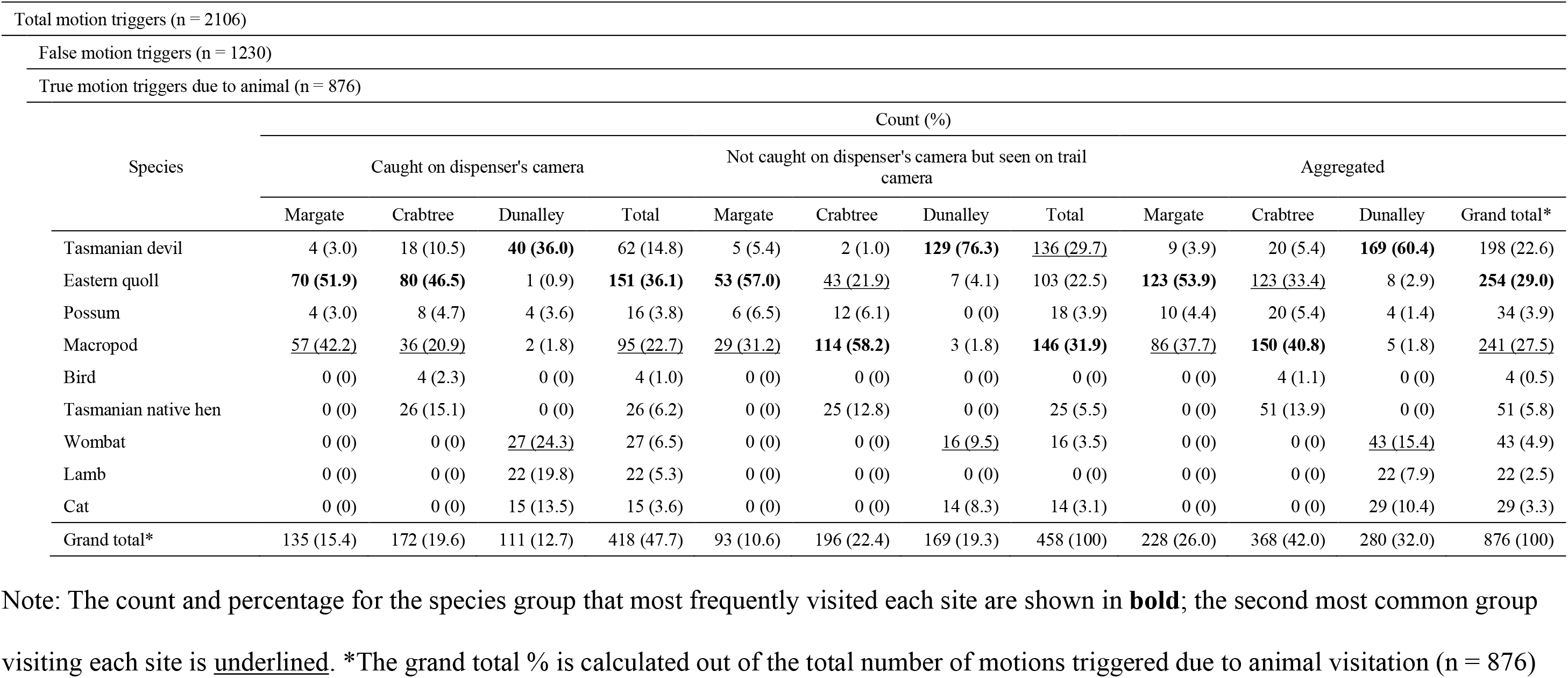
Summary of motion triggers and animal visitations across the three trial sites.

| Total motion triggers (n = 2106) |  |  |  |  |  |  |  |  |  |  |  |  |
| --- | --- | --- | --- | --- | --- | --- | --- | --- | --- | --- | --- | --- |
| False motion triggers (n = 1230) |  |  |  |  |  |  |  |  |  |  |  |  |
| True motion triggers due to animal (n = 876) |  |  |  |  |  |  |  |  |  |  |  |  |
| Species | Count (%) |  |  |  |  |  |  |  |  |  |  |  |
|  | Caught on dispenser's camera |  |  |  | Not caught on dispenser's camera but seen on trail camera |  |  |  | Aggregated |  |  |  |
|  | Margate | Crabtree | Dunalley | Total | Margate | Crabtree | Dunalley | Total | Margate | Crabtree | Dunalley | Grand total* |
| Tasmanian devil | 4 (3.0) | 18 (10.5) | **40 (36.0)** | 62 (14.8) | 5 (5.4) | 2 (1.0) | **129 (76.3)** | <u>136 (29.7)</u> | 9 (3.9) | 20 (5.4) | **169 (60.4)** | 198 (22.6) |
| Eastern quoll | **70 (51.9)** | **80 (46.5)** | 1 (0.9) | **151 (36.1)** | **53 (57.0)** | <u>43 (21.9)</u> | 7 (4.1) | 103 (22.5) | **123 (53.9)** | <u>123 (33.4)</u> | 8 (2.9) | **254 (29.0)** |
| Possum | 4 (3.0) | 8 (4.7) | 4 (3.6) | 16 (3.8) | 6 (6.5) | 12 (6.1) | 0 (0) | 18 (3.9) | 10 (4.4) | 20 (5.4) | 4 (1.4) | 34 (3.9) |
| Macropod | <u>57 (42.2)</u> | <u>36 (20.9)</u> | 2 (1.8) | <u>95 (22.7)</u> | <u>29 (31.2)</u> | **114 (58.2)** | 3 (1.8) | **146 (31.9)** | <u>86 (37.7)</u> | **150 (40.8)** | 5 (1.8) | <u>241 (27.5)</u> |
| Bird | 0 (0) | 4 (2.3) | 0 (0) | 4 (1.0) | 0 (0) | 0 (0) | 0 (0) | 0 (0) | 0 (0) | 4 (1.1) | 0 (0) | 4 (0.5) |
| Tasmanian native hen | 0 (0) | 26 (15.1) | 0 (0) | 26 (6.2) | 0 (0) | 25 (12.8) | 0 (0) | 25 (5.5) | 0 (0) | 51 (13.9) | 0 (0) | 51 (5.8) |
| Wombat | 0 (0) | 0 (0) | <u>27 (24.3)</u> | 27 (6.5) | 0 (0) | 0 (0) | <u>16 (9.5)</u> | 16 (3.5) | 0 (0) | 0 (0) | <u>43 (15.4)</u> | 43 (4.9) |
| Lamb | 0 (0) | 0 (0) | 22 (19.8) | 22 (5.3) | 0 (0) | 0 (0) | 0 (0) | 0 (0) | 0 (0) | 0 (0) | 22 (7.9) | 22 (2.5) |
| Cat | 0 (0) | 0 (0) | 15 (13.5) | 15 (3.6) | 0 (0) | 0 (0) | 14 (8.3) | 14 (3.1) | 0 (0) | 0 (0) | 29 (10.4) | 29 (3.3) |
| Grand total* | 135 (15.4) | 172 (19.6) | 111 (12.7) | 418 (47.7) | 93 (10.6) | 196 (22.4) | 169 (19.3) | 458 (100) | 228 (26.0) | 368 (42.0) | 280 (32.0) | 876 (100) |

Note: The count and percentage for the species group that most frequently visited each site are shown in **bold**; the second most common group
visiting each site is underlined. \*The grand total % is calculated out of the total number of motions triggered due to animal visitation (n = 876)

**Table 2.** Confusion matrix of the bait dispenser’s devil classification model CNN1. F_P_, F_N_, T_P_, and T_N_ denote false positive, false negative, true positive, and true negative, respectively The model was successful in identifying devils in 48 motion triggers and non-devils in 2028 motion triggers.

|  |  | Predicted class |  |
| --- | --- | --- | --- |
|  |  | Devil | Non-devil |
| Actual class | Devil | 48<br>(T<sub>P</sub>) | 14<br>(F<sub>N</sub>) |
|  | Non-devil | 16<br>(F<sub>P</sub>) | 2028<br>(T<sub>N</sub>) |

**Table 3.** Site-wise bait distribution.

|  | Margate | Crabtree | Dunalley | **Total\*** |
| --- | --- | --- | --- | --- |
| True positive | 2 (100) | 11 (57.9) | 18 (90.0) | **31 (75.6)** |
| False positive | 0 (0) | 8 (42.1) | 2 (10.0) | **10 (24.4)** |
| **Total\*** | **2 (4.9)** | **19 (46.3)** | **20 (48.8)** | **41 (100)** |

Note: Numbers indicate the number of baits dispensed and the percentage is in parentheses.
\*The total % is calculated out of the total number of baits dispensed (n = 41)

## Discussion

The smart bait dispenser successfully recognised Tasmanian devils under field conditions, dispensed baits, and facilitated bait consumption, demonstrating the functionality of the system under field settings. The overall bait delivery success rate during the field trials was 75.6%. The system was configured to release a bait when at least five of the ten images captured during a motion event were classified as devil. In several cases, devils appeared in fewer than five images, preventing bait release even when the target species was present. The parameter of requiring five images to be classified as devil and the classification probability threshold of 0.5 were based on early trials. Retraining the model with images clicked during the field trials will allow us to optimise the parameters, i.e., determine presence of a devil using fewer images with greater classification probability. We will also explore alternative robust approaches for inference, like using logistic regression, in future studies.

A key finding was the durability of the bait dispenser housing and design, particularly given that devils routinely crack bones with their powerful jaws. Trail camera footage recorded multiple quolls, possums, and devils attempting to forcibly remove baits from the dispensers. Despite these persistent efforts, no individual could retrieve a bait if the dispenser did not release one in their presence.

We trialled the dispensers in winter conditions and found that they were resilient to rain and frost without compromising the electronic components. However, images collected under inclement weather conditions were of lower quality (**Fig. 5 (a) to (c)**). Modifications, such as a small hood/shield fixed to the top of the dispenser, could minimise the effects of weather conditions and improve image quality. Additionally, many images were not focused on the visiting animal due to its quick movement near the dispenser (**Fig. 5 (d) to (f)**).

Fish oil was effective in drawing animals into the intended field of sight to improve monitoring results (**Fig. 6 (a)**). However, devils, quolls, and possums often moved directly towards the baits inside the dispenser, rather than pausing to investigate the fish oil attractant. Consequently, the animals’ proximity to the dispenser camera meant they were often outside the dispenser camera’s field of view (**Fig. 5 (g) to (h) and Fig. 6 (b) to (f)**). Increasing the concentration or changing the location of fish oil, or trialling other long-distance scent lures could be tested in future studies. Alternative dispenser mounting options could be explored that include physical barriers to steer the visiting animals into the dispenser’s field of view.

The LoRaWAN communication system allowed us to effectively monitor activity at the remote locations without rapid loss of battery power. In addition to receiving exciting activity updates, LoRaWAN communication was useful in identifying if a device required maintenance. For example, the dispenser at Margate stopped operating on the fourth night and this was also indicated by the absence of activity on the online dashboard. This information allowed us to attend the site with a replacement battery and troubleshooting equipment. We found that a software bug in the main program caused the dispenser to fail, hence draining the battery. Since the bug was easy to fix, updating the program and replacing the battery solved the issue. Remotely available activity data could also identify dispensers that require bait replenishment, particularly at sites with high visitation or false triggers. Thus, we conclude that remote monitoring minimises unnecessary site visits and associated labour costs in large-scale deployments.

We achieved an average battery runtime of 19.2 nights across the three trial sites, meeting our objectives for reasonable applications in real world settings. Variations in field conditions like temperature and humidity can affect battery longevity in outdoor applications (reviewed in Shi et al. (2025)). Our trials in Tasmania were conducted during winter, which is expected to reduce battery life due to cold temperatures (Jaguemont et al., 2016). In future studies, we will explore methods to further improve the bait dispenser’s battery life using hardware and software modifications, alternative battery chemistries, or the addition of a small but durable solar panel connection.

The field trials also revealed limitations in our approach to use the smart bait dispenser. False detections are a common limitation of outdoor applications like trail cameras that use passive infrared sensors (Newey et al., 2015). More than half of all motion events were caused by environmental conditions rather than animals. These triggers generated unnecessary images and increased energy consumption, reducing battery life. Methods to minimise environmental noise like alternative mounting methods, sensor angles, and shielding arrangements will be trialled in future studies.

Our smart bait dispenser can distinguish the target species from non-target species but cannot identify individuals. Individual recognition may be important in ecological applications that require controlled dosing. However, highly successful oral rabies vaccine programs distribute baits into the environment at a specified number of baits dropped per square kilometre, resulting in individual animals receiving an unknown and uncontrolled number of vaccine doses. Despite this largely uncontrolled dosing, large-scale oral rabies vaccination drives have been successful (reviewed in Rupprecht et al. (2024)). The computational platform used in our dispenser may support an automated individual-recognition model as technology advances and our image library increases. In the case of devil facial tumour disease, devils with mid to late-stage disease have large facial tumours that can be observed in trail camera images. It may be possible to integrate recognition of facial tumours into the dispenser operational workflow and online dashboard.

In conclusion, we developed a portable and adaptable smart bait dispenser that holds up to 20 baits, runs on a rechargeable battery pack, and uses a simple image classification model to identify the target species within seconds. The cost to manufacture this smart bait dispenser is currently estimated at US$350 per unit based on costs in 2024-2025. The Arduino Portenta H7 Vision Shield (LoRa) was the most expensive component, accounting for approximately one-third of the total device cost contributor (approximately US$210). Given our objective of developing an open-source design, this dispenser could be modified by users to lower production costs and support further technological development. We made our bait dispenser open-source so that anyone can adapt it for any target species by retraining the onboard machine learning model. The dispenser proved suitable for multi-day field deployments and remained operational under Tasmanian weather conditions. These findings highlight the significant potential of automated, species-selective bait dispensers to advance wildlife conservation efforts. Our versatile system can be adapted for supplementary feeding, delivering preventative or therapeutic medicines, and precise target-selective trapping.

## Acknowledgements

We acknowledge the Palawa/Tasmanian Aboriginal people as the traditional owners of Lutriwita/Tasmania. We thank Eagle Plastics and Phil Hortin for professionally cutting PVC pipes for the dispensers. We thank the Arducam Active Users program for supplying Himax HM0360 NoIR cameras. We thank Adrian Hamilton, David Chung, Eddy Roccati, Ibrahim, Abdelkader, Jeremy Ellis, Jon Nordby, Joshua Eastgate, Kwabena Agyeman, Marcelo Rovai, Marco Zennaro, Peter Padd, Soyeon Caren Han, Indicium Dynamics, and the online community of Arduino, Micropython, Stack Overflow and Youtube for their contributions in troubleshooting software issues. We thank Wynne Russell and Felicity Smart (Margate), Jenny Cambers-Smith (Crabtree), and Matt Dunbabin (Dunalley) for access to their private properties and allowing us to organise the field trials. We thank Alexandre Kreiss, Anuk Kruawan, Emma Nguyen, Grace Russell, Jocelyn Darby, Lauren Snell, Macarena Gonzalez Abarzua, Mary Fisher, Ruth Pye, Sandesh Sharma, Thomas Walsh, and the staff members of Bonorong Wildlife Sanctuary and the Department of Natural Resources and Environment Tasmania for their support in bench and field testing the dispenser. We thank Animal Control Technologies for provision of placebo baits, and Gregory Woods for his feedback on the manuscript. This study was funded by the Australian Research Council grants LP210301148, LP230200956, and FT240100092 awarded to Andrew S. Flies. The study was also supported by funds raised through the Save the Tasmanian Devil Appeal, donations by Tim and Sheena High, and Roger and Maxeme Tall through the Tall Foundation, Pure Foods Eggs, AAT Kings, and many smaller donors.

## Author contributions

Prithul Chaturvedi, Andrew S. Flies, and William M. Connelly conceptualised the smart bait dispenser. Prithul Chaturvedi, Andrew S. Flies, William M. Connelly, and Tim Lennard led the hardware development. Prithul Chaturvedi and William M. Connelly led the software development. Prithul Chaturvedi, Andrew S. Flies, Chrissie Ee Been Ong, and Timothy J. Smyser planned and executed the field trials. Prithul Chaturvedi and Andrew S. Flies led the writing of the manuscript. All authors contributed critically to the drafts and gave final approval for publication.

## Data availability statement

Design files and programs used in this manuscript are archived on [Zenodo](https://zenodo.org/records/23383887?preview=1&token=eyJhbGciOiJIUzUxMiJ9.eyJpZCI6IjJlMTdmMTBhLTNhZWEtNDgyOS05MzcwLTZmMjQxYjQwYWQwZCIsImRhdGEiOnt9LCJyYW5kb20iOiI0YjE3MTE4YzRjOTQxNGZkYTU4MTQyYmJjOTJlNTQ5MiJ9.E-woQ5GCgKvf-vUDfxTwQ97abgoI57SBLGhZTEZYvce-vDl3I71Du6ArUICqDY8Mv4DYMK3ozd6HIVD7KjnPfg). Latest project documentation and resources are available at: https://github.com/prithulc/smart-bait-dispenser.

