## Supplementary material for "A Smart Bait Dispenser with Real-time Monitoring for Species-Specific Bait Delivery": Figure S1

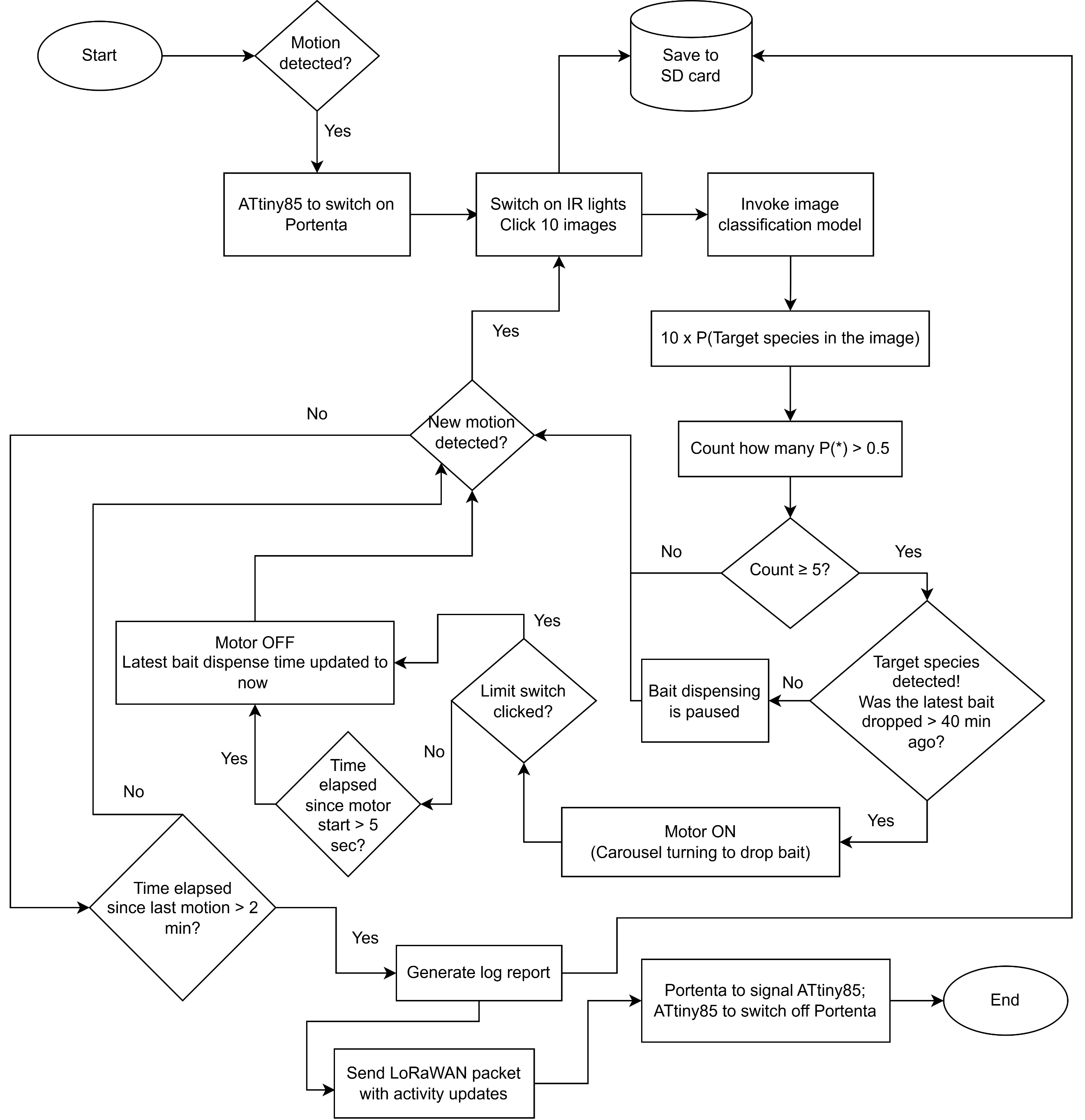

**Figure S1** Detailed flowchart of the bait dispenser’s operational workflow process. Made with drawio (Jgraph, 2021).
